## Supplementary Information for "Facilitating Analysis and Dissemination of Proteomics data through Metadata Integration in MaxQuant"

### Step-by-step protocol for exporting the SDRF file in MaxQuant, and using SDRF to annotate output tables in Perseus

**These features have been implemented since MaxQuant 2.7.0 and Perseus 2.1.5. The video tutorial can be viewed on YouTube at [https://youtu.be/fHCPOBXXRp8](https://youtu.be/fHCPOBXXRp8).**

### 1. Set up the MaxQuant run

1.1 Load the raw files and set experiment (also set the fractions if you have them)

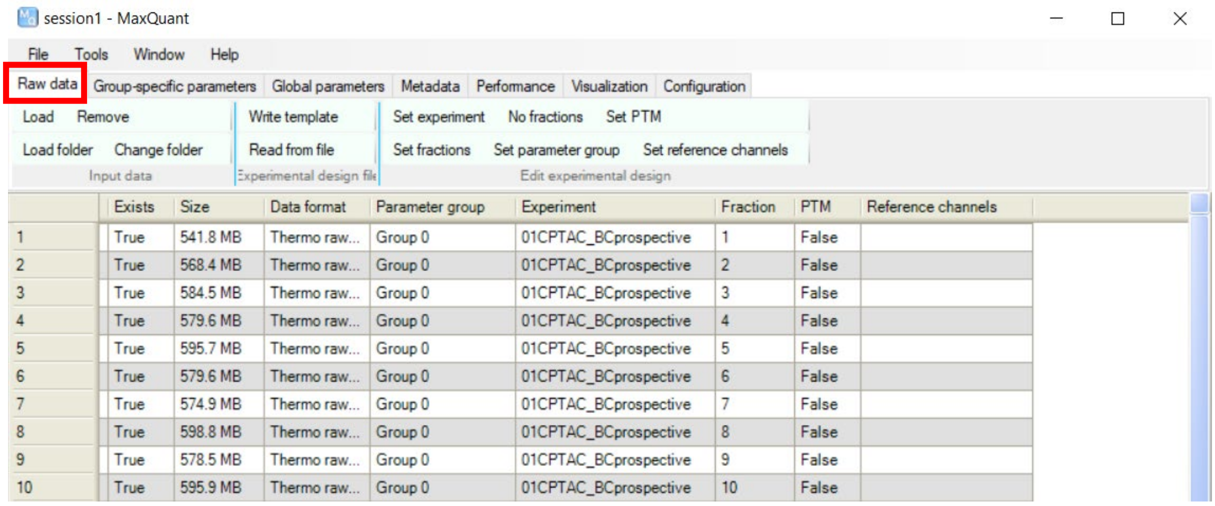

1.2 Set the Group-specific parameters for the run

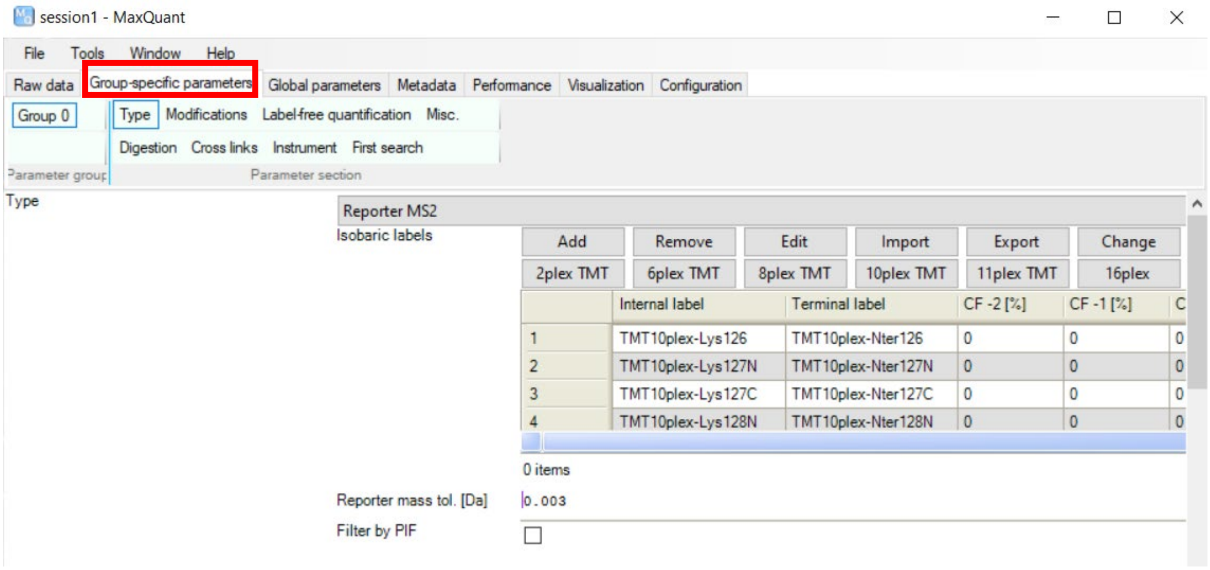


1.3 Set the Global parameters for the run

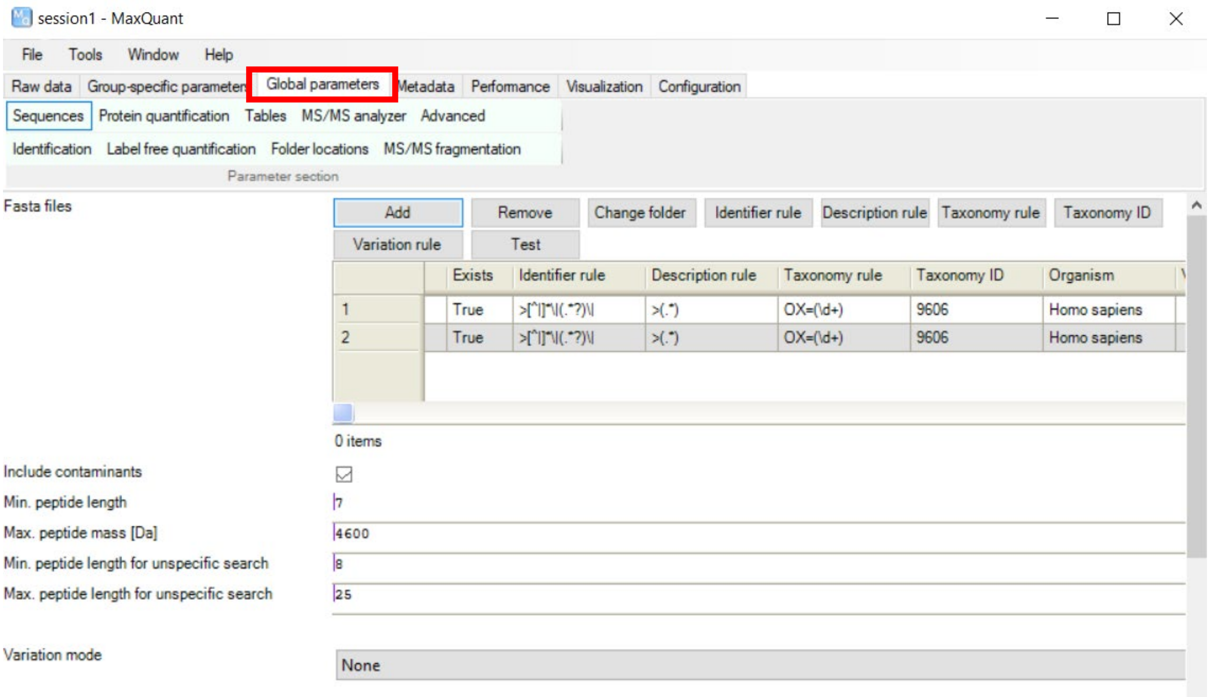

### 2. Export SDRF in MaxQuant

2.1 Create the metadata table by click “Refresh” under the “Metadata” tab

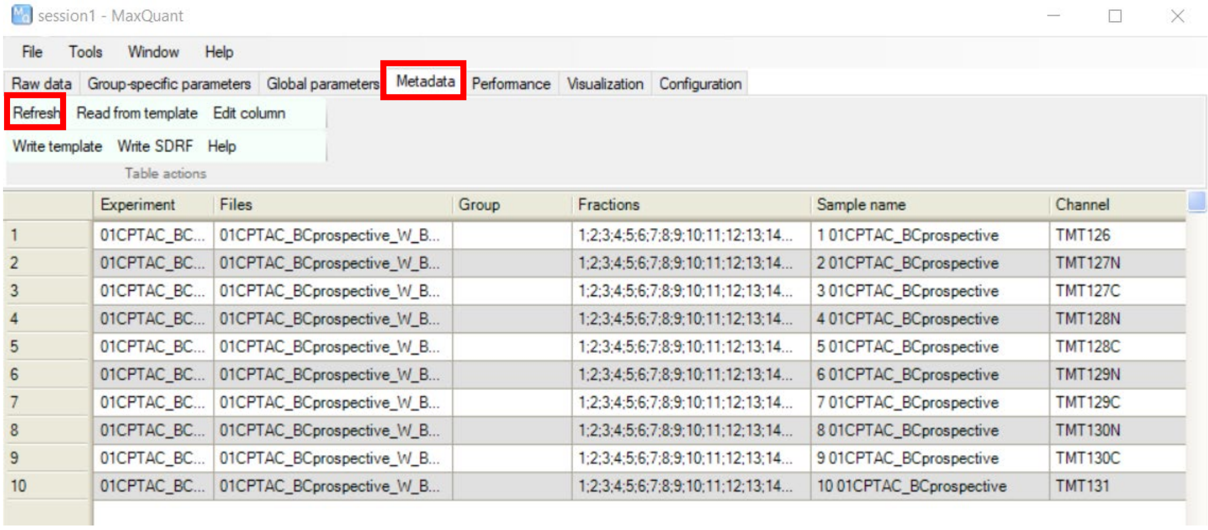


2.2 Select the column(s) and edit column(s) in GUI. If information for certain columns are not available in your study, leave them empty. MaxQuant will fill the empty cells with “not available” when writing SDRF

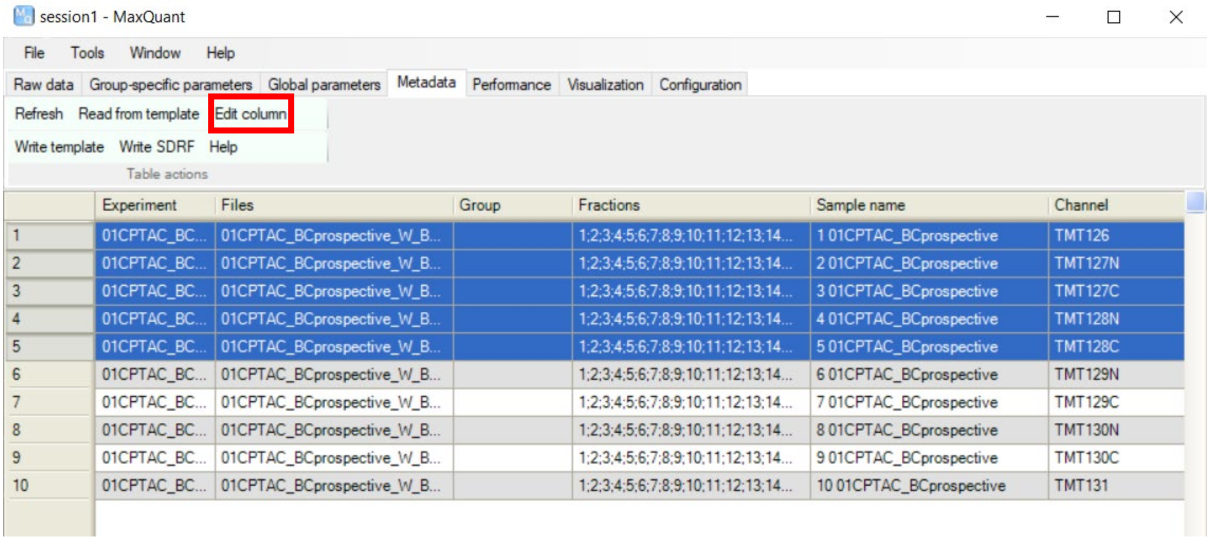

Or write template for metadata table, fill it and read from the filled template

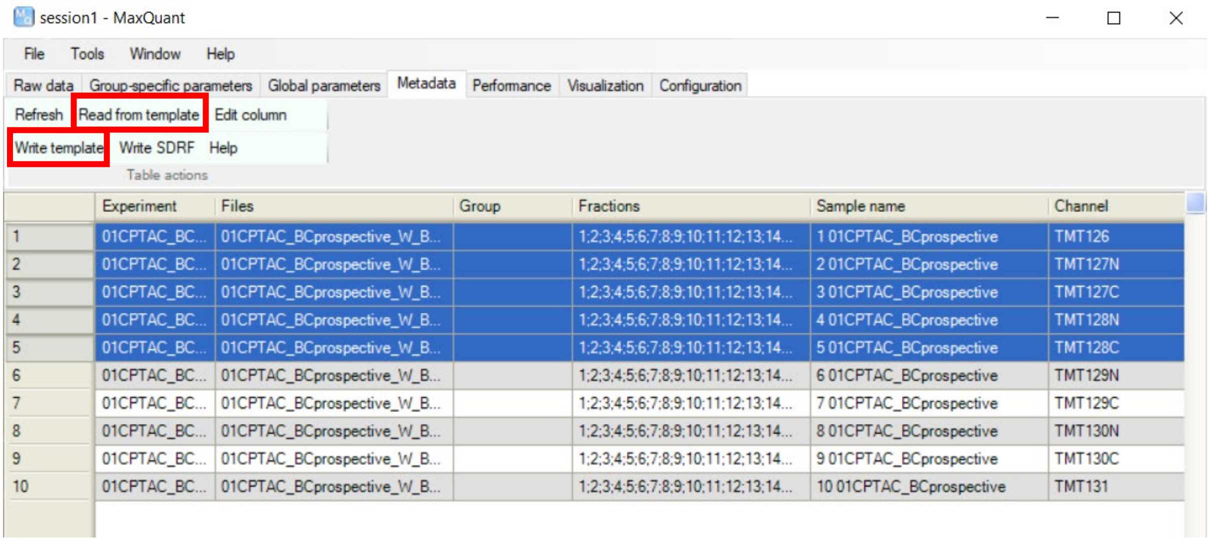


| | Experiment | Files | Group | Fractions | Sample name | Channel |
|---|---|---|---|---|---|---|
| 1 | 01CPTAC_BC... | 01CPTAC_BCprospective_W_B... | | 1;2;3;4;5;6;7;8;9;10;11;12;13;14... | 1 01CPTAC_BCprospective | TMT126 |
| 2 | 01CPTAC_BC... | 01CPTAC_BCprospective_W_B... | | 1;2;3;4;5;6;7;8;9;10;11;12;13;14... | 2 01CPTAC_BCprospective | TMT127N |
| 3 | 01CPTAC_BC... | 01CPTAC_BCprospective_W_B... | | 1;2;3;4;5;6;7;8;9;10;11;12;13;14... | 3 01CPTAC_BCprospective | TMT127C |
| 4 | 01CPTAC_BC... | 01CPTAC_BCprospective_W_B... | | 1;2;3;4;5;6;7;8;9;10;11;12;13;14... | 4 01CPTAC_BCprospective | TMT128N |
| 5 | 01CPTAC_BC... | 01CPTAC_BCprospective_W_B... | | 1;2;3;4;5;6;7;8;9;10;11;12;13;14... | 5 01CPTAC_BCprospective | TMT128C |
| 6 | 01CPTAC_BC... | 01CPTAC_BCprospective_W_B... | | 1;2;3;4;5;6;7;8;9;10;11;12;13;14... | 6 01CPTAC_BCprospective | TMT129N |
| 7 | 01CPTAC_BC... | 01CPTAC_BCprospective_W_B... | | 1;2;3;4;5;6;7;8;9;10;11;12;13;14... | 7 01CPTAC_BCprospective | TMT129C |
| 8 | 01CPTAC_BC... | 01CPTAC_BCprospective_W_B... | | 1;2;3;4;5;6;7;8;9;10;11;12;13;14... | 8 01CPTAC_BCprospective | TMT130N |
| 9 | 01CPTAC_BC... | 01CPTAC_BCprospective_W_B... | | 1;2;3;4;5;6;7;8;9;10;11;12;13;14... | 9 01CPTAC_BCprospective | TMT130C |
| 10 | 01CPTAC_BC... | 01CPTAC_BCprospective_W_B... | | 1;2;3;4;5;6;7;8;9;10;11;12;13;14... | 10 01CPTAC_BCprospective | TMT131 |

2.3 Click “Write SDRF” to write the SDRF file. The SDRF file will also be generated automatically once the MaxQuant run is complete

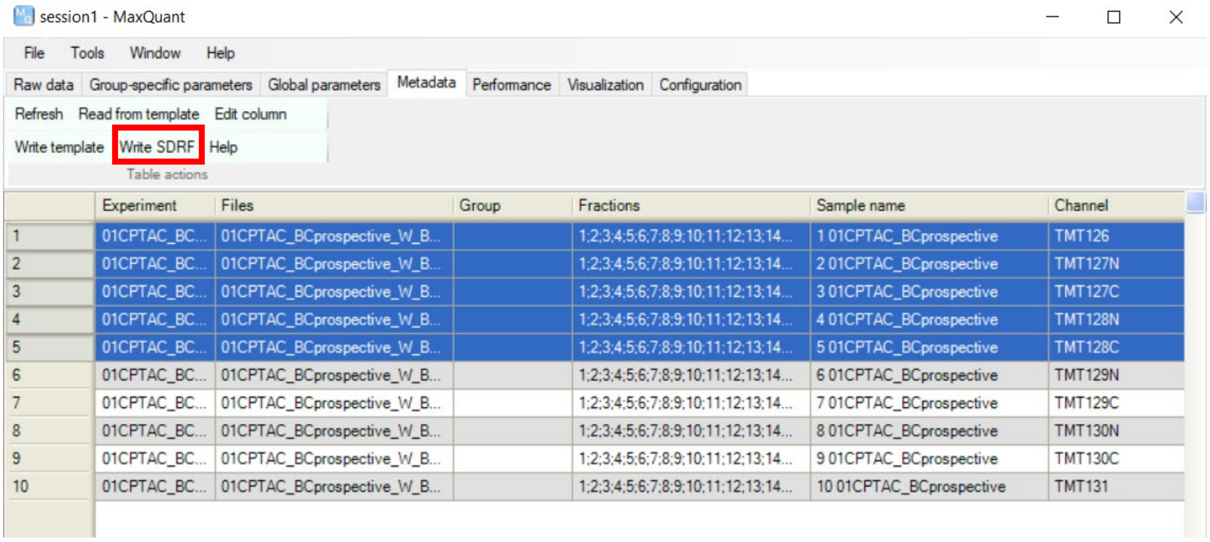


| | Experiment | Files | Group | Fractions | Sample name | Channel |
|---|---|---|---|---|---|---|
| 1 | 01CPTAC_BC... | 01CPTAC_BCprospective_W_B... | | 1;2;3;4;5;6;7;8;9;10;11;12;13;14... | 1 01CPTAC_BCprospective | TMT126 |
| 2 | 01CPTAC_BC... | 01CPTAC_BCprospective_W_B... | | 1;2;3;4;5;6;7;8;9;10;11;12;13;14... | 2 01CPTAC_BCprospective | TMT127N |
| 3 | 01CPTAC_BC... | 01CPTAC_BCprospective_W_B... | | 1;2;3;4;5;6;7;8;9;10;11;12;13;14... | 3 01CPTAC_BCprospective | TMT127C |
| 4 | 01CPTAC_BC... | 01CPTAC_BCprospective_W_B... | | 1;2;3;4;5;6;7;8;9;10;11;12;13;14... | 4 01CPTAC_BCprospective | TMT128N |
| 5 | 01CPTAC_BC... | 01CPTAC_BCprospective_W_B... | | 1;2;3;4;5;6;7;8;9;10;11;12;13;14... | 5 01CPTAC_BCprospective | TMT128C |
| 6 | 01CPTAC_BC... | 01CPTAC_BCprospective_W_B... | | 1;2;3;4;5;6;7;8;9;10;11;12;13;14... | 6 01CPTAC_BCprospective | TMT129N |
| 7 | 01CPTAC_BC... | 01CPTAC_BCprospective_W_B... | | 1;2;3;4;5;6;7;8;9;10;11;12;13;14... | 7 01CPTAC_BCprospective | TMT129C |
| 8 | 01CPTAC_BC... | 01CPTAC_BCprospective_W_B... | | 1;2;3;4;5;6;7;8;9;10;11;12;13;14... | 8 01CPTAC_BCprospective | TMT130N |
| 9 | 01CPTAC_BC... | 01CPTAC_BCprospective_W_B... | | 1;2;3;4;5;6;7;8;9;10;11;12;13;14... | 9 01CPTAC_BCprospective | TMT130C |
| 10 | 01CPTAC_BC... | 01CPTAC_BCprospective_W_B... | | 1;2;3;4;5;6;7;8;9;10;11;12;13;14... | 10 01CPTAC_BCprospective | TMT131 |

### 3. Annotate output tables with SDRF file in Perseus

3.1 Load the output tables in Perseus (peptides.txt, proteinGroups.txt, etc.), Specify the intensity columns as "Main"

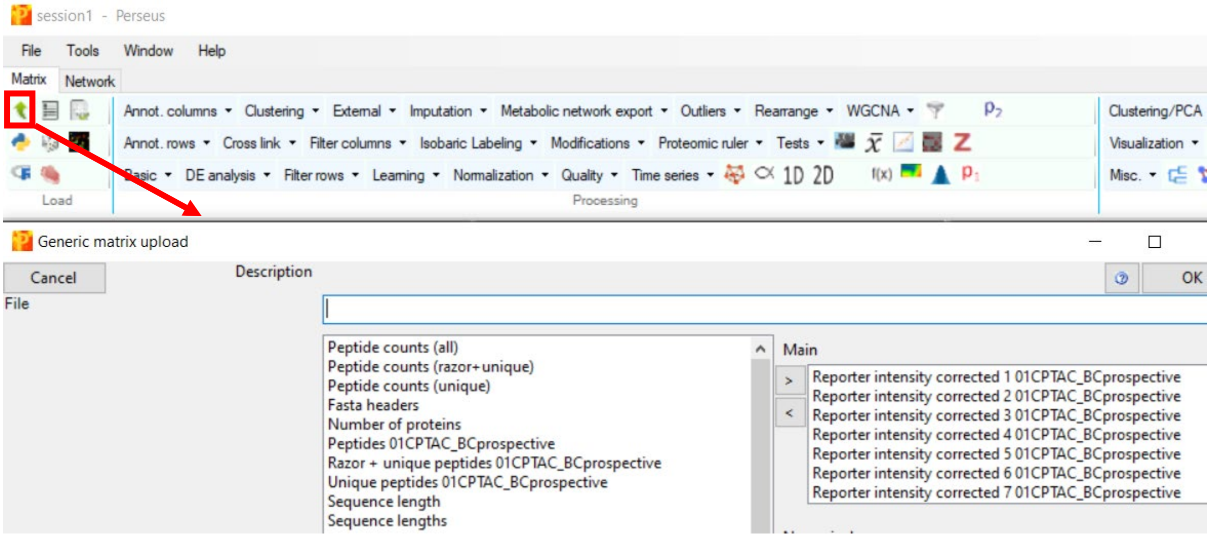


3.2 Read SDRF for the annotation, by default "Skip redundant properties" is selected

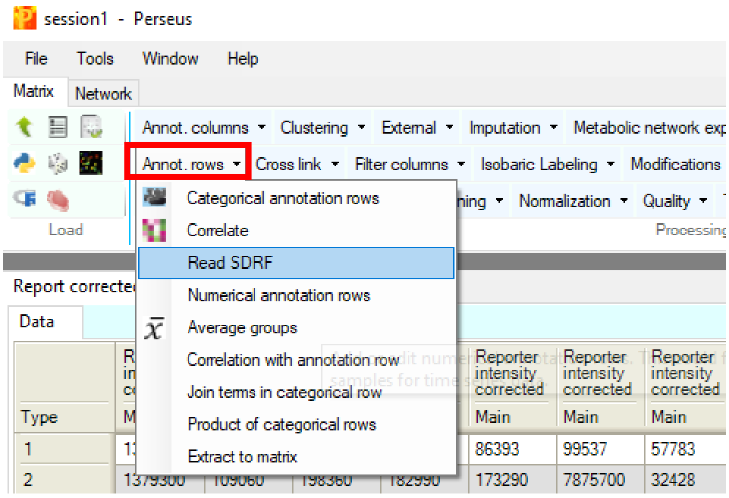


Alternatively, Python and R scripts are provided (https://github.com/cox-labs/converters) to convert MaxQuant output tables and the SDRF file into an expression matrix and a sample annotation table, which are required by other popular tools for downstream data analysis, such as DESeq2, limma, and edgeR.
